# Pharmacological and fasting-induced activation of SIRT1/LXRα signaling alleviates diabetes-induced retinopathy

**DOI:** 10.1101/871822

**Authors:** Sandra S. Hammer, Cristiano P. Vieira, Delaney McFarland, Maximilian Sandler, Yan Levitsky, Tim F. Dorweiler, Todd Lydic, Bright Asare-Bediako, Yvonne Adu-Agyeiwaah, Micheli Severo Sielski, Mariana Dupont, Ana Leda Longhini, Sergio Li Calzi, Dibyendu Chakraborty, Gail M. Seigel, Denis A. Proshlyakov, Maria B. Grant, Julia V. Busik

## Abstract

In diabetes, the retina, a tissue with unique metabolic needs, demonstrates dysregulation of the intricate balance between nutrient availability and utilization. This results in cholesterol accumulation, pro-inflammatory and pro-apoptotic changes, and consequently neurovascular damage. Sirtuin 1 (SIRT1), a nutrient sensing deacetylase, is downregulated in the diabetic retina. In this study, the effect of SIRT1 stimulation by fasting or by pharmacological activation using SRT1720, was evaluated on retinal cholesterol metabolism, inflammation and neurovascular damage. SIRT1 activation, in retinal endothelial cells (REC) and neuronal retinal progenitor cells (R28), led to Liver X Receptor alpha (LXRα) deacetylation and subsequent increased activity, as measured by increased ATP-binding cassette transporter (ABC) A1 and G1 mRNA expression. In turn, increased cholesterol export resulted in decreased REC cholesterol levels. SIRT1 activation also led to decreased inflammation. SIRT1 activation, *in vivo*, prevented diabetes-induced inflammation and vascular and neural degeneration. Diabetes-induced visual function impairment, as measured by electroretinogram and optokinetic response, was significantly improved as a result of SIRT1 activation. Taken together, activation of SIRT1 signaling is an effective therapeutic strategy that provides a mechanistic link between the advantageous effects associated with fasting regimes and prevention of diabetic retinopathy (DR).

## Introduction

The high metabolic demands of the retina dictate complex regulatory pathways needed to meet retinal energy needs while preserving autonomy behind the blood-retina barrier (BRB). Healthy retinal function is maintained by a delicate balance between nutrient availability and tightly regulated retinal lipid metabolism (1). However, in a diabetic environment, characterized by chronic low-grade inflammation and lipid accumulation, the balance between nutrient availability and retinal-specific metabolism may be lost (2-4). One of the major consequences of dysregulation of lipid metabolism is elevated retinal cholesterol levels, ultimately fostering the progression of DR, the most common microvascular complication (5, 6).

Previously we, and others, have demonstrated the advantageous effects of activation of cholesterol regulating mechanisms in the prevention of DR progression. Specifically, we have shown that activation of LXRα in the diabetic retina, by synthetic ligand administration, stimulates the reverse cholesterol transport (RCT) pathway resulting in decreased retinal cholesterol accumulation (7, 8). Additionally, activation of the LXRα signaling cascade reduces diabetes-induced inflammation by repressing the NF-kβ response element on inflammatory genes such as IL-1β and IL-6 (7). Consequently, due its ability to regulate retinal cholesterol levels and anti-inflammatory properties, activation of LXRα signaling effectively prevents DR progression *in vivo*. However, activation of LXRα by pharmacological ligands, have the potential to cause a compensatory increase in expression of SREBP1c and ChREBP resulting in hypertriglyceridemia and liver steatosis (9, 10). Thus, additional efforts for identifying alternative mechanisms of LXRα activation are warranted.

In addition to ligand binding, LXRα can be activated via deacetylation. SIRT1 deacetylase is a major controller of LXR acetylation status (11). We have previously shown that pharmacological SIRT1 activation, *in vitro*, results in elevation of LXRα mediated signaling (7). SIRT1 is a NAD^+^-dependent nutrient sensing deacetylase and is activated in nutrient scarce conditions (12, 13). SIRT1 is uniquely positioned to sense a tissues metabolic status and fine tune tissue demands to maintain a homeostatic balance.

Due to the importance of SIRT1-mediated signaling in metabolism, deregulation of SIRT1 deacetylase activity results in chronic metabolic abnormalities. For instance, in diabetes, SIRT1 expression and activity are significantly decreased (14, 15). In the diabetic retina, activation of SIRT1 is protective against diabetes-induced vascular and mitochondrial damage by inhibiting MMP-9 activation (14). Importantly, diabetes-induced decreases in SIRT1 signaling results in dysregulated cholesterol metabolism and increased production of pro-inflammatory cytokines via decreased LXRα signaling (7). However, *in vitro* under diabetes mimicking conditions, pharmacological activation of SIRT1 prevents increases in cholesterol accumulation and inhibits inflammation by activating LXRα signaling (7).

Due to SIRTs dependence on NAD^+^, a strong association has been made between SIRT activation and periods of low nutrient availability. Fasting regimes and SIRT1 activation have been shown to increase longevity and delay onset of disease in yeast, fruit flies, mice and more recently, non-human primate animal models (12, 16, 17). Fasting-induced SIRT1 activation has been linked to increased NAD^+^ levels, increased mitochondrial biogenesis and delayed senescence (17-19). Additionally, we showed that chronic intermittent fasting prevents the development of DR and extends longevity by restructuring the microbiome of type 2 diabetic animals (20). The mechanisms regulating these beneficial outcomes include the increased generation of endogenous oxysterols and their activation of the nuclear receptor TGR-5 within the retina; however additional mechanisms are under active investigation.

The current study was designed to determine the role of the SIRT1/LXRα signaling pathway in prevention of DR. Pharmacological agents were used to activate SIRT1 both in *in vitro* and *in viv*o models as well as fasting mimicking conditions (FMCs) *in vitro*. SIRT1/ LXRα signaling was examined by measuring RCT activation and inhibition of NF-Kβ-dependent proinflammatory gene production. Thus, modulation of nutrient availability and subsequent SIRT1/LXR activation, provides an innovative mechanism to prevent DR.

## Results

### Diabetes-induced retinal cholesterol accumulation and inflammation is prevented by SIRT1 activation

The diabetic retinal milieu is characterized by increased levels of inflammation and cholesterol accumulation. Moreover, SIRT1/LXR signaling is significantly decreased in diabetes-associated complications, including DR (7, 21). In order to investigate the effect SIRT1 activation on DR progression, control and *db/db* animals were fed chow containing the selective SIRT1 activator, SRT1720 (100mg/kg body weight/day) for 6 months following the onset of diabetes. Diabetes significantly reduced SIRT1 retinal expression. However, retina isolated from mice fed SRT1720-containing chow had increased retinal SIRT1 expression, with preservation of the ganglion cell layer (GCL) when compared to retinas from diabetic mice fed control chow (**Figure 1A-D**).

**Figure 1.**
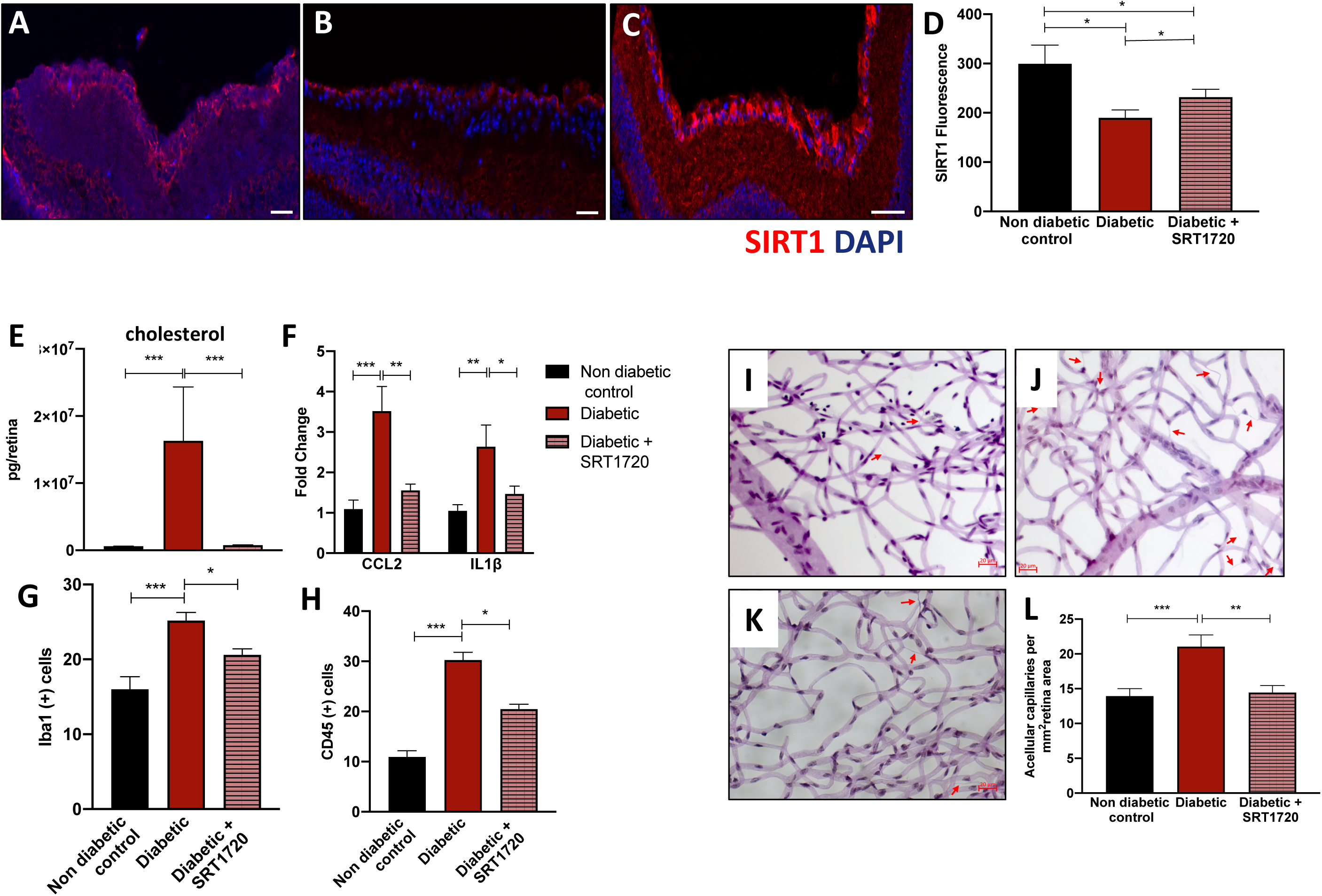
SRT1720 increases SIRT1 expression in retina of *db/db* mice with 6-month duration of diabetes. Retinal sections from (**A**) non-diabetic controls, diabetic mice fed (**B**) normal chow or (**C**) chow containing SRT1720 were stained with anti-SIRT1 antibody (red). Diabetes significantly decreases SIRT1 expression in the ganglion cell layer (GCL) while pharmacological activation using SRT 1720 prevents diabetes-induced SIRT1 downregulation. Quantification is shown in **D**. DAPI used to stain nuclei (blue). n=4, **E**) Diabetes significantly increases retinal cholesterol levels, the SIRT1 agonist SRT 1720 restores cholesterol levels to non-diabetic levels. Diabetes significantly increases pro-inflammatory markers (**F**) CCL-2, IL-1β and increases (**G**) Iba1(^+^), (**H**) CD45(^+^) cells. n=5, (**I-L**) Acellular capillary formation (red arrows) was examined in (**I**) non-diabetic, and diabetic animals fed (**J**) control chow or (**K**) chow containing STR1720. Diabetes significantly increases acellular capillary formation (**J**) while administration of the SIRT1 agonist prevents diabetes-induced acellular capillary formation (**K**). Quantification shown in **L**. n=5 *p<0.01, **p<0.001, ***p<0.0001. Data are represented as mean ± SEM. Scale bar: 50μm.

SIRT1 has been previously shown to deacetylate, and subsequently, activate LXR. Increased SIRT1/LXR signaling led to increased levels of RCT, via upregulation of ABCA1 and ABCG1, and upregulation of anti-inflammatory gene expression (7). As expected, diabetes resulted in elevated retinal cholesterol levels when compared to control retinas. SIRT1 activation, via SRT1720, restored cholesterol levels in diabetic retina to non-diabetic levels (**Figure 1E**). Treatment with SRT1720 also alleviated diabetes-induced inflammation, as measured by expression of IL-1β, CCL-2 and Iba-1^+^ and CD45^+^cells (**Figure 1F-H**). Moreover, activation of SIRT1 signaling, prevented diabetes-induced downregulation of ABCG1 expression in anti-inflammatory M2 macrophages (supplemental material S1A).

### SIRT1 activation prevents retinal vascular and neurodegeneration *in vivo*

Formation of acellular capillaries is the hallmark feature of DR. As shown in **Figure 1I**, the development of acellular capillaries was significantly increased in *db/db* mice (red arrows). Activation of SIRT1, via SRT1720 treatment, restored the number of acellular capillaries to non-diabetic levels (**Figure 1K-L**). Neuronal damage and loss (**Figure 2**) precede the microangiopathy of DR. As expected, diabetes resulted in a loss of neurons as measured by NeuN^+^ immunohistochemistry analysis (**Figure 2B**). However, administration of SRT1720 significantly preserved NeuN^+^ expression, predominately localized in the retinal ganglion cell layer (**Figure 2C**).

**Figure 2.**
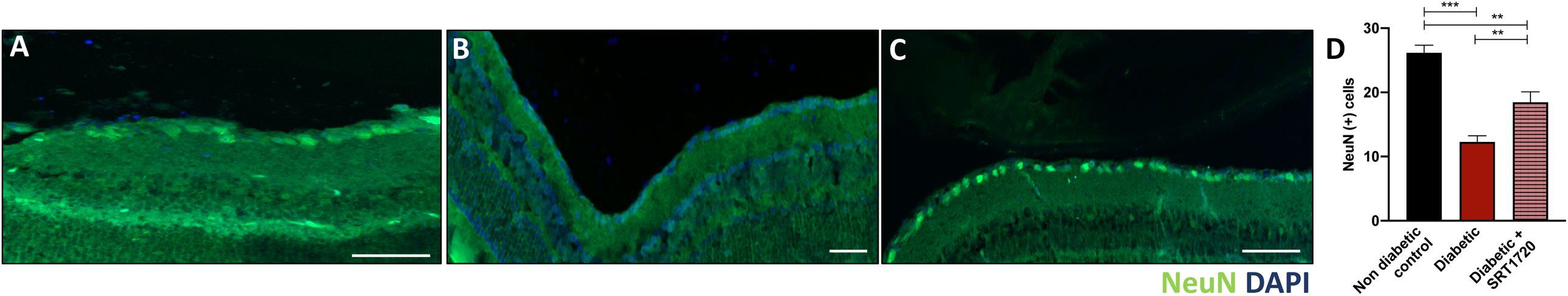
SRT1720 prevents diabetes-induced NeuN^+^ retinal decrease. Chronic diabetes is associated with neuronal loss in the retina as demonstrated by reduced NeuN^+^ expression (green) in *db/db* mice (**B**) when compared to (**A**) non-diabetic db/m controls. **C**) Pharmacological SIRT1 activation using STR1720 in *db/db* mice partly restores NeuN^+^ expression to control levels. Quantification shown in **D**. n=4,**p<0.001, ***p<0.0001. Data are represented as mean ± SEM. Scale bar: 50μm

### Fasting-mimicking conditions activate SIRT1/LXR signaling in vascular cells and neuronal cells

SIRT1 is activated by low nutrient, calorie reduced physiological states. In addition to pharmacological activation of SIRT1, via SRT1720, fasting mimicking conditions (FMCs) (0% FBS) activated SIRT1 expression and histone deacetylase (HDAC) activity in REC (**Figure 3A**). Consequently, increased HDAC activity resulted in elevated total LXRα protein levels and activity, as measured by expression of ATP-binding cassette transporter A1 and G1 ABCA1,ABCG1; respectively)(**Figure 3B**). Moreover, total and active LXRα protein levels were significantly increased in REC treated with FMC (**Figure 3D-F**).

**Figure 3.**
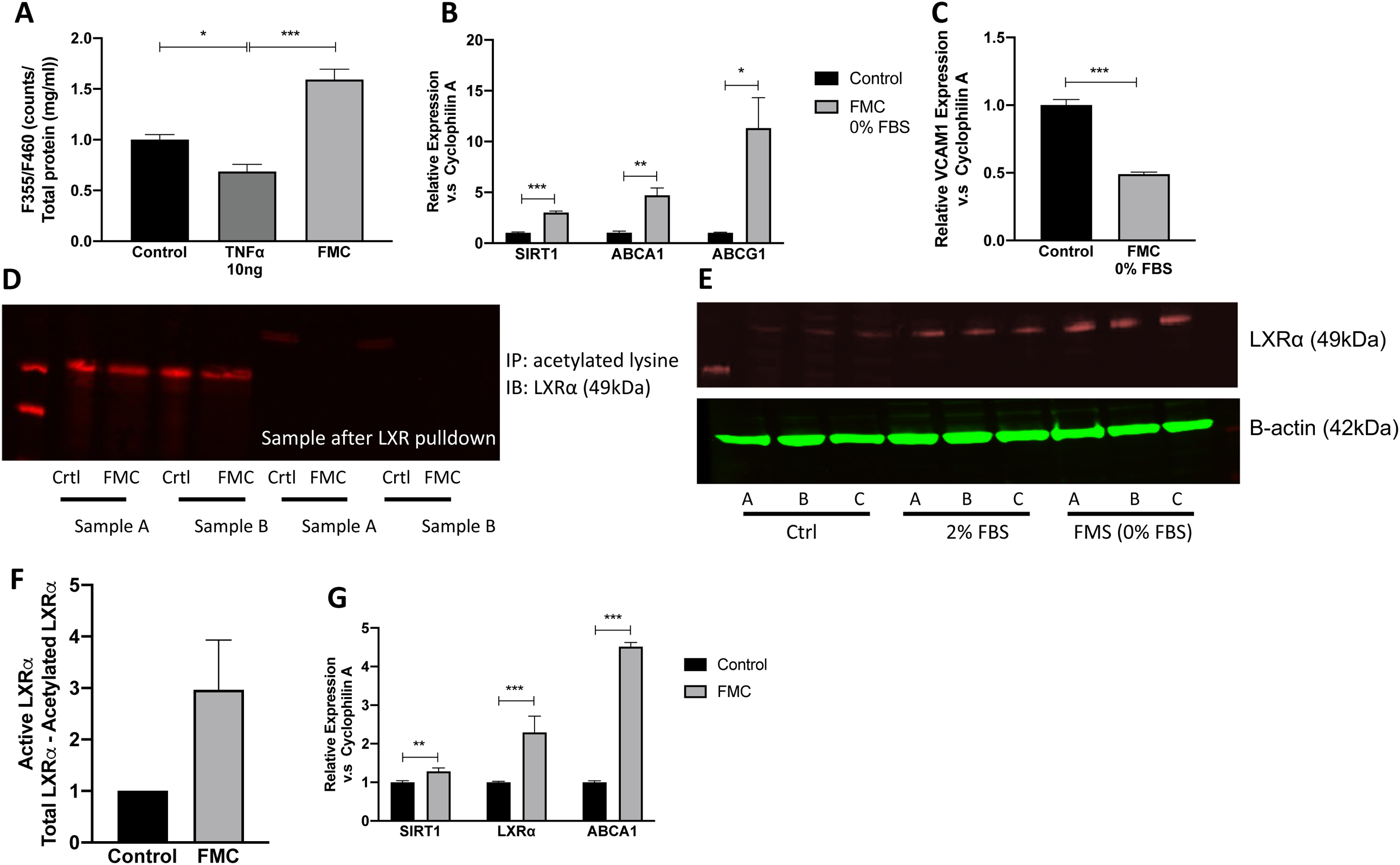
Fasting mimicking conditions (FMC) activate the SIRT1/LXR signaling pathway in retinal endothelial (REC) and neuronal (R28) cells. Fasting mimicking conditions (FMC; 0% FBS) in REC activate (**A**) the HDAC activity of SIRT and (**B**) increased reverse cholesterol export (ABCA1 and ABCG1). **C**) Inflammation (VCAM1) is decreased after treatment with FMCs (24hrs). FMC treatment (**D**) decreases non active, deacetylated LXR levels and **E**) increases total LXRα protein levels. Active LXRα levels, (acetylated lysine-total control and 0% FBS) are quantified in **F**. Crtl (10% FBS), FMC (0% FBS). **G**) Treatment of R28 cells with FMC (0% FBS, 24hrs) increases expression of SIRT1 mRNA and LXR activity (ABCA1). n=6; *p<0.01, **p<0.001, ***p<0.0001. Data are represented as mean ± SEM.

As shown in **Figure 4A**, REC administrated TNFα which is typically increased in diabetes, resulted in increased cholesterol levels. In contrast, FMC lowered cholesterol accumulation in REC (supplemental material S1B). As expected, activation of LXRα via DMHCA, a steroidal LXR ligand, significantly reduced REC cholesterol levels. Treatment with FMC, in combination with DMHCA administration, significantly augmented cholesterol export and lowered REC cholesterol levels even further when compared to DMHCA treatment alone (**Figure 4B**).

**Figure 4.**
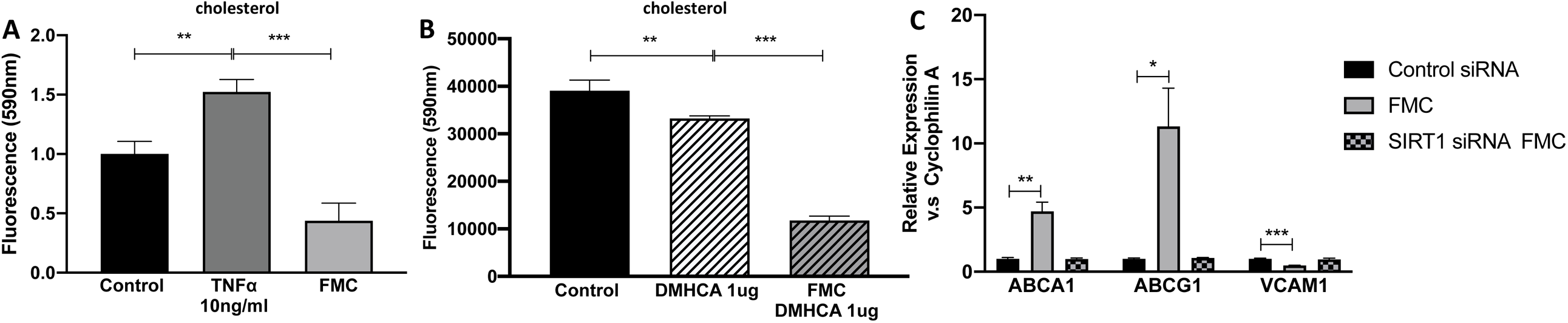
SIRT1 plays an important role in fasting mediated decrease levels of cholesterol in retinal endothelial cells (REC). **A**) TNFα treatment causes a significant increase in cholesterol levels while FMC (24hrs) results in decreased levels of cholesterol. This decrease is amplified with (**B**) administration of the LXRα activator, DMHCA. **C**) SIRT1 was reduced by exposure of the REC to SIRT1 siRNA (80% knockdown efficiency, data not shown). The administration of SIRT siRNA prevented serum-induced upregulation of ABCA1 and ABCG1 and inhibited the downregulation of the pro-inflammatory gene, VCAM1. n=6; **p<0.001, ***p<0.0001. Data are represented as mean ± SEM.

In addition to augmenting cholesterol export, FMC in REC prevented the upregulation of the NF-kβ dependent pro-inflammatory gene, VCAM1 (**Figure 3C**). Importantly, data shown in **Figure 4** highlights the importance of SIRT1 signaling in regulating the FMC-induction of RCT and the anti-inflammatory responses in REC. Loss of SIRT1, via SIRT1-directed siRNA, prevents FMC-induced upregulation of ABCA1, ABCG1 and VCAM1 (**Figure 4C**).

As previously demonstrated in **Figure 2**, pharmacological activation of SIRT1 prevents diabetes-induced neurodegeneration. In order to assess the effect of FMC-induced SIRT1 activation in neuronal cells, SIRT1/LXR activity was measured in rat retinal progenitor cells (R28 cells). Treatment with FMC significantly increased SIRT1 and LXRα expression as well as caused an increased in LXRα activity, as measured by ABCA1 expression (**Figure 3G**).

### Cell death and mitochondrial respiration are not affected by FMC

Prolonged reduced serum nutrient conditions have the potential to induce cell death or cause mitochondrial damage. In order to access the possible cytotoxic effects of FMC treatment, cell death and mitochondrial respiration were measured after 24hrs in FMC-treated RECs. As shown in **Figure 5A** cell death, measured by Trypan Blue exclusion assay, did not significantly differ between FMC-treated cells and REC cultured in control conditions (10% FBS containing medium) (supplemental material S1C). Moreover, there is no statistical difference between mitochondrial respiration in cells cultured in FMC or control conditions (**Figure 5B**, Supplemental material S2).

**Figure 5.**
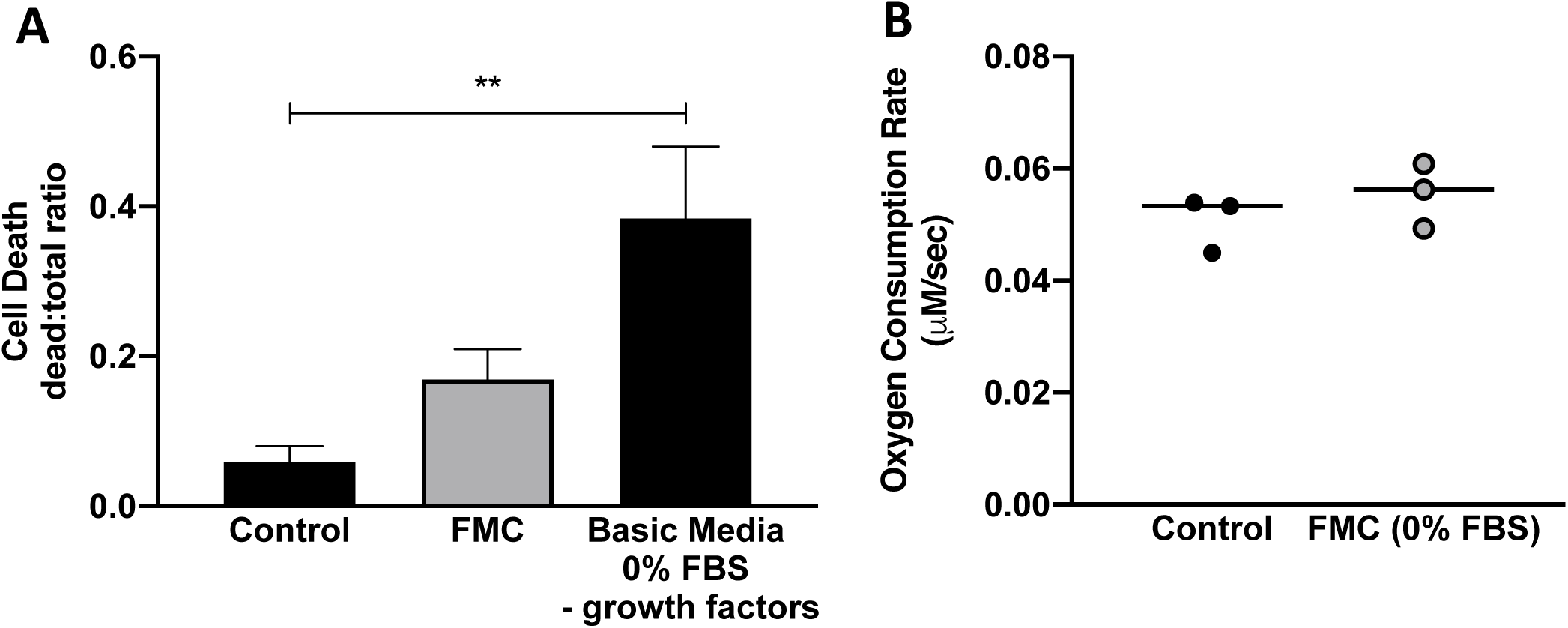
Fasting mimicking conditions (FMC) do not significantly impact cell death or mitochondrial substrate supported respiration in retinal endothelial cells (REC). FMC (0% FBS; 24hrs) does not significantly increase **A**) cell death as measured by Trypan blue exclusion assay. **B**) After 24 hours of perfusion with serum free media, the respiratory activity was 55.5 ± 5.8 nM/sec (n = 3) compared to from 50.7 ± 5.0 nM/sec n= 3). **p<0.001, Data are represented as mean ± SEM.

### SIRT1 activation improves visual function in *db/db* mice

Histopathological neurovascular changes in the diabetic retina result in functional deficiencies including electroretinogram (ERG) and visual response abnormalities. The functional effect of SIRT1 activation in diabetes was assayed by measuring retinal and visual function of *db/db* mice fed control or SRT1720-containing chow. As expected, ERG of *db/db* mice showed decreased scotopic a, b waves and lower photopic b waves. ERG responses were restored to non-diabetic levels in SIRT1 1720 treated diabetic mice (**Figure 6A-B**). Visual response assayed using optokinetic response (OKN) measurements were significantly lower in diabetic mice when compared to control mice and SIRT1 activation restored visual acuity to control levels (**Figure 6C-D**).

**Figure 6.**
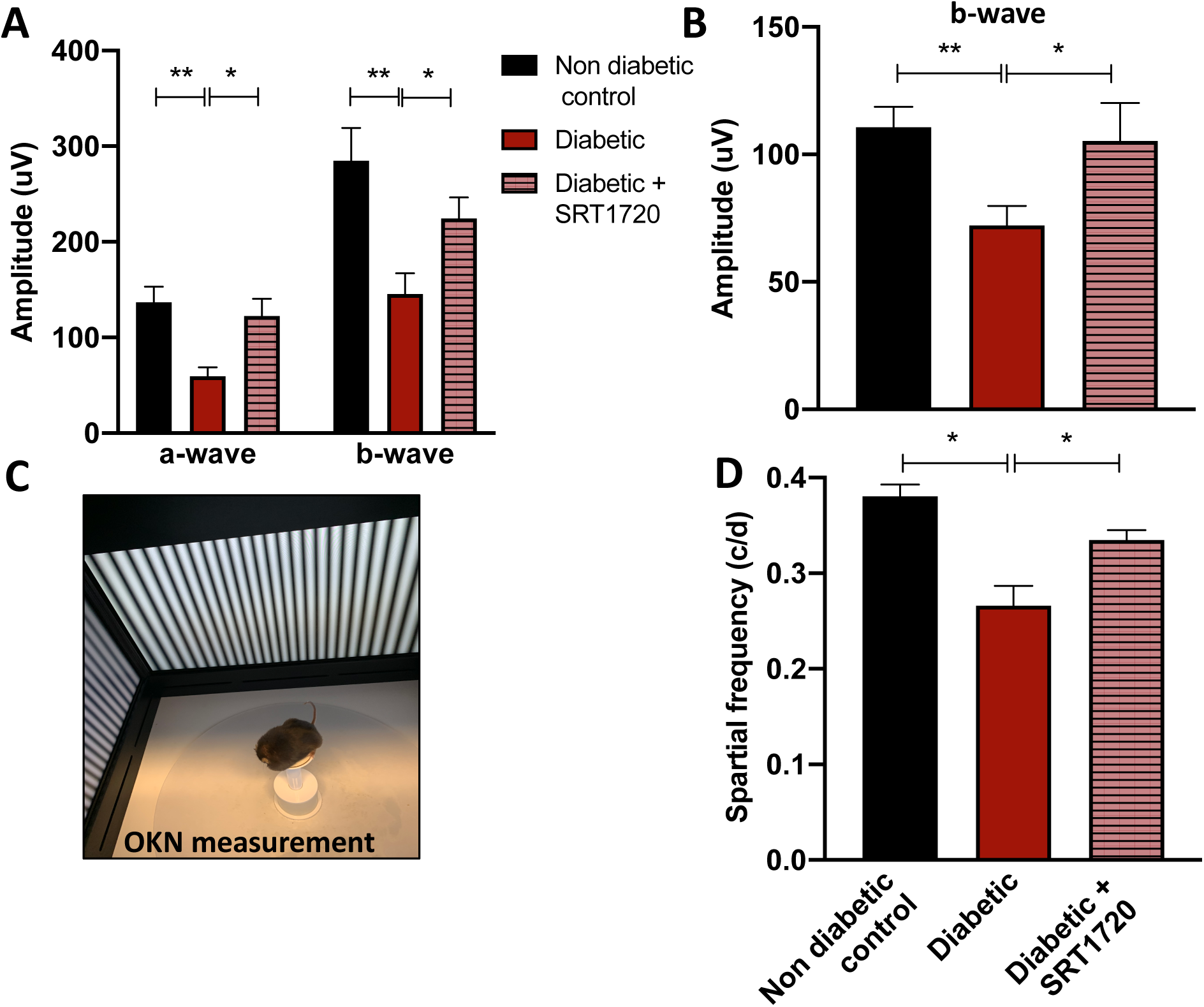
Activation of SIRT1 improves visual and optokinetic response in *db/db* mice. Analysis of the electroretinogram (ERG) showed diabetes induced reduction of scotopic a-and b-waves. An improvement in scotopic (**A**) a- and b-wave was observed in *db/db* mice treated with SRT1720 compared to *db/db* mice on control chow. **B**) An improvement in photopic b-wave was observed in *db/db* mice treated with SRT1720 compared to *db/db* mice on control chow but no difference was observed in the photopic a-wave. n=5, **C-D**) In *db/db* mouse, the OKN response is reduced compared to age matched non-diabetic db/m mice. Diabetic *db/db* mice treated with SRT1720 showed an improvement in visual acuity of the compared with *db/db* (diabetes) mice. n=6,*p<0.05 *p<0.01, **p<0.001, ***p<0.0001. 1 Data are represented as mean ± SEM.

## Discussion

High demand for cholesterol in the retina dictates its unique metabolism with local retinal cholesterol production coupled with import from the circulation. Under healthy conditions cholesterol enters the retina via the outer blood retinal barrier through low density lipoprotein receptor (LDL) mediated processes (22).

Rapid cholesterol turnover in the retina requires efficient cholesterol export. Retinal cells express several hydroxylases (CYP 27A1, CYP 46A1, CYP 11A1) that metabolize cholesterol to more soluble oxysterols (23). Unlike cholesterol, oxysterols can rapidly diffuse out of the retina for the delivery to the liver for generation of bile acids. Additionally, oxysterols produced by CYPs 27A1 and 46A1 activate LXR, promoting LXR-mediated retinal cholesterol export through activation of RCT pathway via increased ABCA1 and ABCG1 expression in both inner (REC) and outer (RPE) retinal barrier.

In the diabetic environment, both cholesterol uptake and removal from the retina are affected, resulting in elevated and dysregulated retinal cholesterol levels. The leaky retinal endothelial cells lead to increased cholesterol ingress, and the decrease in oxysterol levels and LXR activity result in cholesterol accumulation in the retina.

In addition to the increase in RCT, activation of LXRα results in repression of NF-Kβ dependent pro-inflammatory gene upregulation in the retinal cells. Low LXR expression and activity levels contribute to pro-inflammatory conditions in the diabetic retina. Activation of LXR is thus an attractive target for normalizing cholesterol metabolism and inflammation under diabetic conditions. Indeed, we have previously shown that activation of LXRα via steroidal synthetic LXRα ligand (GW 3965 and DMHCA) halts progression of DR by regulating levels of cholesterol in the retina and preventing diabetes-induced inflammation (7, 8).

Direct LXRα activation, however, increases expression of SREBP1c and ChREBP leading to liver steatosis, highlighting the need for innovative approaches for modulation of LXR activity (9, 10). In the current study, we utilized SIRT1 nutrient sensing abilities to deacetylate and activate LXRα and its downstream signaling targets. This work demonstrates an innovative and effective way of activating the SIRT1/LXR signaling pathway and subsequent prevention of retinal metabolic disease. Figure 1 demonstrates the detrimental effects type 2 diabetes has on retinal SIRT1 expression, cholesterol metabolism and inflammation. In diabetic mice, activation of SIRT1 via SRT1720 restores SIRT1 expression and cholesterol levels to normal. Additionally, diabetes-induced retinal inflammation is significantly reduced (**Figure 1F-H**). This increase in SIRT1 expression results in decreased inflammation and increased cholesterol export (**Figure 1E**). Additionally, activation of SIRT1 has beneficial physiological effects as seen in the prevention of DR-induced acellular capillary formation (**Figure 1I-L**), a hallmark of DR progression. Notably, *in vivo* activation of SIRT1 restores visual function in *db/db* mice as measured by ERG and OKN response (**Figures 6**).

SRT1720 has been previously shown to increase lifespan and improve overall health in mice fed a high fat diet as well as those fed a normal diet (24, 25). Activation of SIRT1, via SRT1720, has been shown to significantly improve insulin sensitivity, lower glucose levels and increase mitochondrial and metabolic function (26). SRT1720 was shown to activate SIRT1 with potencies 1,000-fold greater than resveratrol, another SIRT1 activator (26). Although administration of SRT1720 was successful in activating retinal SIRT1 levels in our studies, future work on potential off target effects of SRT1720 treatment is needed. Other groups have shown that SIRT1720 is not a direct activator of SIRT1 in non-retinal tissue. Additionally, Pacholec et. al. suggests that activation of SIRT1 via SRT1720 does not lower plasma glucose nor improve mitochondrial capacity in high fat diet fed mice (27). In contrast, others have shown SRT1720 as a potential therapeutic agent for type 2 diabetic individuals (26). There is contradictory evidence as to the pro-survival properties of SIRT1 activation. Zarse et al. demonstrated the negative effects that SRT1720 has on C. elegans lifespan (28). However, this group also showed that treatment with resveratrol increased C. elegans lifespan, suggesting that in this specific model organism, the effects seen by SIRT1 activation via resveratrol cannot be duplicated by SRT1720. These confounding effects could be due to the governing role that SIRT1 plays in regulating circadian rhythm in peripheral tissues. Several studies have demonstrated SIRT1 activation of BMAL1 and CLOCK genes, resulting in robust circadian restoration (29, 30). Thus, differences in timing and feeding patterns among different studies could result in varied physiological outcomes.

In addition to pharmacological activation of SIRT1, we show that FMCs activate SIRT1/LXRα signaling in retinal endothelial and neuronal cells (**Figure 3**). This finding is particularly exciting due to the numerous studies highlighting the beneficial effects intermittent fasting and calorie restriction regimes have on preventing diabetes-associated complications (20, 31-33). Recently, the beneficial effects of calorie restriction were highlighted by a clinical trial in which subjects who maintained reduced caloric intake for two years had a significant reduction in heart disease and diabetes prevalence (34). However, the mechanism of action responsible for these beneficial effects are still under active investigation.

In sum, the work highlighted in this manuscript demonstrates the beneficial effects of fasting or pharmacological activation of SIRT1/LXRα signaling on retinal RCT, inflammation and retinal function in the diabetic milieu. Importantly, this study positively contributes to the growing evidence for the effective and advantageous effects of modulating time and amount of nutrient intake.

## Acknowledgements

This study was supported by the National Institutes of Health Grants R01EY012601, R01EY028858, R01EY028037, R01EY025383 to M.B. Grant; T32HL134640-01 to M. Dupont; T32HL105349 to J.L. Floyd, F32EY028426 to S.S. Hammer; MICL02539, R01EY016077 to J.V.Busik, R01EY028049 to D.A. Proshlyakov; NIH-5-R25-HL108864 to M.Sandler. Dr. Marina Gorbatyuk and Prof. Timothy W. Kraft (School of Optometry, UAB) provides LKC for ERG and OptoMotry for visual acuity study respectively.

## Author Contributions

Conceptualization- S.S.H, M.B.G., J.V.B., C.P.V.

Methodology- S.S.H, M.B.G., J.V.B, D.A.P. C.P.V.; A.L.L.

Investigation- S.S.H., C.P.V.,D.M.,M.S.,Y.L.,T.F.M.,T.L., Y.A.A.; M.S.S.; M.D., A.L.L., S.L.C.; D.C.

Writing – Original Draft, S.S.H and J.V.B.

Writing – Review & Editing, S.S.H., M.B.G., J.V.B, C.P.V. D.M, M.S., T.F.D.,T.L. D.A.P, Y.L.

Visualization – S.S.H., C.P.V; M.B.G and J.B.V.

Project administration- C.P.V, M.B.G.; S.S.H.

Funding Acquisition- J.V.B., M.B.G., D.A.P., S.S.H.

Resources- J.V.B.,M.B.G., G.M.S.

Supervision- J.V.B., M.B.G.

## STAR Methods

### Animal studies

All animal procedures were in compliance with the National Institutes of Health (NIH) *Guide for the Care and Use of Laboratory Animals*, and with the Association for Research in Vision and Ophthalmology *Statement for the Use of Animals in Ophthalmic and Vision Research* (IACUC # 10917). Mouse eyes were enucleated, and retinas were harvested by “Winkling” as described by Winkler and placed in RNALater storage solution (35). For *db/db* (Strain (B6.BKS(D)-*Lepr*^db^/J (Stock#000697) was purchased from Jackson Laboratory) studies, mice were treated with 100mg/kg body weight/day SRT1720 in their chow for 6 months after the diabetic onset around 10 weeks of mice age (36). Mice were considered diabetic after two measurements of 250 mg/dL of glucose level. *db/db* and db/m without treatment were analyzed as control animals. Retinas were analyzed for qRT-PCR and lipid quantification. Immunofluorescence studies were performed using frozen sections for NeuN^+^, Iba1^+^, CD45^+^ cells and SIRT1 expression. Acellular capillaries were verified by trypsin digestion. The visual response was verified by ERG and scotopic a-wave and b-wave were detected as well as the photopic b-wave response. Optokinetic response was performed to detection of visual motion on the retina.

### Cell culture and treatment

Bovine retinas were isolated from eyes donated by the Michigan State University Meat Laboratory. Bovine retinal endothelial cells (BREC) were isolated and validated according to a previously published protocol (37). BREC identity was confirmed morphologically and using anti-Von Willebrand Factor immunocytochemistry (Abcam Cat# ab6994, RRID:AB_305689) and LDL uptake assay (ab133127) (38). BREC were cultured in 10% Fetal Bovine Serum (FBS) BREC Complete Media with 1% Antibiotic/Antimycotic (AA) (Gibco; ThermoFisher; Waltham, MA). Passages 4–8 were used for all experiments. In order to model a fasting environment *in vitro*, BREC were cultured in 0% FBS for 24hrs (fasting mimic conditions, FMC). FMC media contained endothelial growth factors and 5mM glucose. The heterogeneous adherent rat retinal neuroglial/precursor R28 cell line was acquired from Gail M. Seigel at the University at Buffalo. R28 cells were cultured in DMEM+ media with 10% calf serum. Cells were treated with diabetic relevant stimuli tumor necrosis factor alpha for 24 h (10 ng/ml) (2279-BT; R&D Systems) (39).

### Cell death assay

BREC were cultured in FMC and after the treatment period cell death was analyzed via Trypan Blue Exclusion Assay (40). Cell death was calculated by diving the number of dead cells over the total number of cells.

### qRT-PCR

RNA was isolated according with the RNeasy mini kit (74106; Qiagen, Valencia, CA) according to manufacturer’s instructions. First-strand complementary DNA was synthesized from isolated RNA using SuperScript II reverse transcription (18064014; Invitrogen). Prepared cDNA was mixed with 2 × SYBR Green PCR Master Mix (4309155; Applied Biosystems) and various sets of gene-specific forward and reverse primers (SIRT1, ABCA1, ABCG1, VCAM1) (Integrated DNA Technologies; Coralville, IA)) (**Table 1**) and subjected to real-time PCR quantification using the ABI PRISM 7700 Sequence Detection System (Applied Biosystems). All reactions were performed in triplicate. Cyclophilin A was used as a control, and results were analyzed using the comparative Ct method and Ct values were normalized to Cyclophilin A levels. Data is shown as normalized relative to control levels or as non-normalized raw expression levels.

**Table 1.**
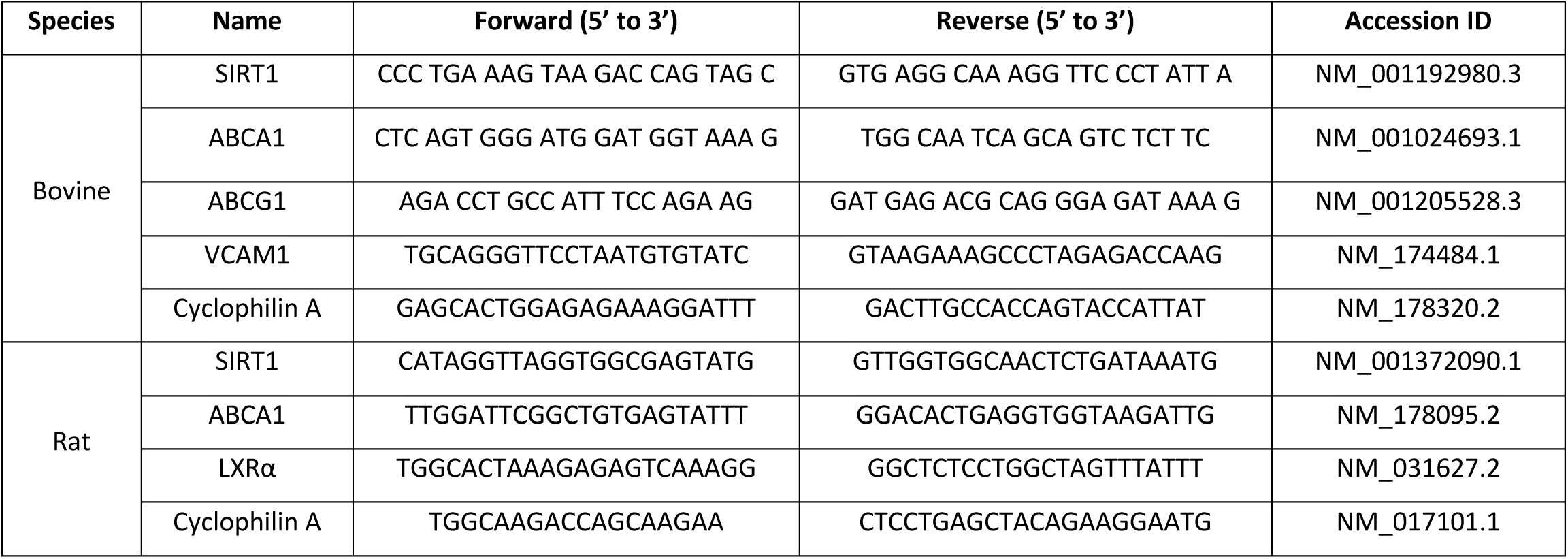
Primer sequences used in qRTPCR analysis.

### Western Blot and IP

BREC were cultured and treated as necessary. Lysates were collected by scraping cells using cold 1 × radioimmunoprecipitation assay buffer with protease and phosphatase inhibitor (89900; Thermo Scientific). Bradford protein assay was used to measure total protein level and equal amounts of protein (25 μg) were loaded in each well. 10% Bis-Tris NuPAGE gels were transferred to 0.45 μm nitrocellulose membranes. Membranes were probed overnight with LXRα (Abcam Cat# ab3585, RRID:AB_303930), and β-actin antibodies (Cell Signaling Technology Cat# 4970, RRID:AB_2223172). β-actin was used as a loading control. Blots were analyzed using the LI-COR Odyssey imaging and quantification system. For immunoprecipitation studies, acetylated lysines were assayed using the Immunoprecipitation Kit Dynabeads Protein A (Invitrogen; 10006D).

### HDAC assay

SIRT1 activity assay was measured using the Histone Deacetylase Activity Assay Kit (ABCAM; ab156064) according to the manufacturer’s instructions. Briefly, 300,000 cells were used for each condition. HDAC assay buffer and substrate was added to each sample and incubated for 45 minutes. Fluorescence intensity was measured at Ex/Em 355/460nm. Total protein was used as a normalization for each sample (mg/ml).

### Mitochondrial respiration measurements

Respiration of BREC’s was measured using an open-shell microrespirometer as previously described (41). Briefly, BREC at passage 4 were seeded and cultured on fibronectin-coated chips overnight at 37° C, 95% RH, 5% CO_2_. On the day of measurements, the chips were closed, mounted in a microrespirometer, and immediately perfused at a constant 10 µL/min flow rate. Perfusion was controlled by a Harvard Apparatus syringe infusion pump. Cells were first perfused with 10% FBS media for 12 hours, followed by perfusion with serum-free media for an additional 24 hours. Respiratory activity was measured for 15 minutes in stationary medium, with a minimum of 15 minutes of re-perfusion between measurements. Total oxygen concentration was maintained above 200 µM in all cases. Measurements in the presence of potassium cyanide (KCN, 5 µM) were used to correct for non-mitochondrial oxygen consumption. Respiratory activity was calculated by linear fitting of the initial five minutes of steady oxygen consumption. Instrument was calibrated used air-equilibrated media as an aerobic standard and 1 mM sodium dithionite in media as anaerobic standard.

### Cholesterol measurement

#### In vivo

##### Lipid extraction

Lipids from mouse retinas were extracted with methanol, chloroform, and water as previously described (42). Prior to tissue homogenization, each retina was spiked with 100 nanograms of 19-hydroxycholesterol (Cayman Chemical, Ann Arbor, MI) for quantitation of sterols. Retina lipid extracts were subjected to alkaline hydrolysis of sterol esters for analysis of total sterol content according to (43).

##### Analysis of Free and Total Sterol Content

Sterols were analyzed by high resolution/accurate mass LC-MS/MS using a Thermo Scientific LTQ-Orbitrap Velos LC-MS system. Gradient conditions, peak finding, and quantitation were performed as previously described (44). Cholesterol identification was performed by comparison of retention time, exact mass, and MS

##### In vitro

Total cholesterol levels were measured using the Amplex Red Cholesterol Assay kit (ThermoFisher Scientific; A12216) according to the manufacturer’s instructions. Briefly, cells were diluted in 1X reaction buffer and incubated with a working solution containing 300uM Amplex Red reagent. Fluorescence was measured at 590nM after 45 minutes at 37°C.

### Immunohistochemical staining

Retinas were removed from eyes of euthanized mice with 6 months duration of diabetes. Retinas were embedded in OCT medium and retinal sections (12μm thickness) were prepared for immunohistochemistry. For the staining process, slides were placed in a solution of (1:1) methanol and hydrogen peroxide for 5 minutes and rinsed once with PBS for 5 min. It was then placed in 5% blocking reagent (goat serum) in PBS for 1hour 30 minutes. Sections were then incubated overnight at 4°C with primary monoclonal antibody for CD45 (R and D Systems Cat# MAB114, RRID:AB_357485) and Iba-1 (Wako Cat# 019-19741, RRID:AB_839504) at ratios of 1:50 and 1:500 respectively. Following this primary incubation, the slides were washed 3X with PBS 1X. Slides were counterstained with Alexa-fluor secondary antibodies. The slides were then washed with PBS X3. DAPI solution was then applied for 10 min after which the slides were washed with PBS and vectashield was applied.

### Visual function studies

#### Electroretinogram (ERG)

For full-field ERG recordings, mice were dark adapted for 12 h. In preparation for the ERGs, the mice were anesthetized with intraperitoneal (IP) injections of ketamine and xylazine. The pupils were dilated with 1% atropine sulfate and 2.5% phenylephrine hydrochloride ophthalmic solution which also reduced sensitivity of the eyes to touch. When fully anesthetized, the mice were placed on a stand in a LED Ganzfeld stimulator (LKC Technologies, Gaithersburg, MD). A drop of Goniotaire solution (from Altaire pharmaceuticals and contains 2.5% hypromellose solution) was applied to each eye to ensure a good electrical connection between the electrodes and the eyes. Animals were kept on a warm (37°C) heating pad during the procedure. Full-field ERGs were recorded from both eyes using the LKC system. Corneal electrodes with contact lenses were placed on each eye, and a steel subdermal needle served as the reference electrode. For grounding, a steel needle was placed in the tail.

To test scotopic responses, animals received a series of flashes at intensities −20db, −10db and 0db. For photopic responses, flashes were at intensities −3, 3, 6 and 10db after 5 minutes of light adaptation. The inter-stimulus intervals were longer and with fewer trials at higher intensities to reduce bleaching of rod photopigments (45).

#### Optomotor response

The visual acuity of the mice was determined by use of optomotor response recordings (46, 47). This was quantified from the spatial frequency as measured by a trained observer-operator. The presentation of stimuli from computer monitors displaying vertical dark and white lines in motion elicits visual tracking reflex in the mice that are being tested. The test was performed using an optomotor system (OptoMotry; Cerebral Mechanics, Inc., Lethbridge, Canada). An individual mouse was placed on a platform inside the chamber of the system. The mouse moves the head reflexively as the vertical stripes move in the chamber with the direction of motion of the head following the movement of the stripes to show tracking behavior. The system automatically varies the spatial frequency of the stripes depending on whether the mouse successfully tracked the stripes or not and arrives at a final spatial frequency. This is the highest spatial frequency recorded for the mouse. This test was done under photopic conditions with the contrast between the gratings at 100% and speed of 12.0 d/s.

### Statistics

A Student *t-*test was used for comparisons between two groups. One-way ANOVA, followed by the Tukey post hoc test, was used for multiple comparisons. All values are expressed as mean ± SEM. A value of p < 0.05 was considered to be statistically significant. Statistical tests were performed using statistics software (GraphPad Software; La Jolla, CA).

## Supplemental Information

**Supplemental Figure 1.**
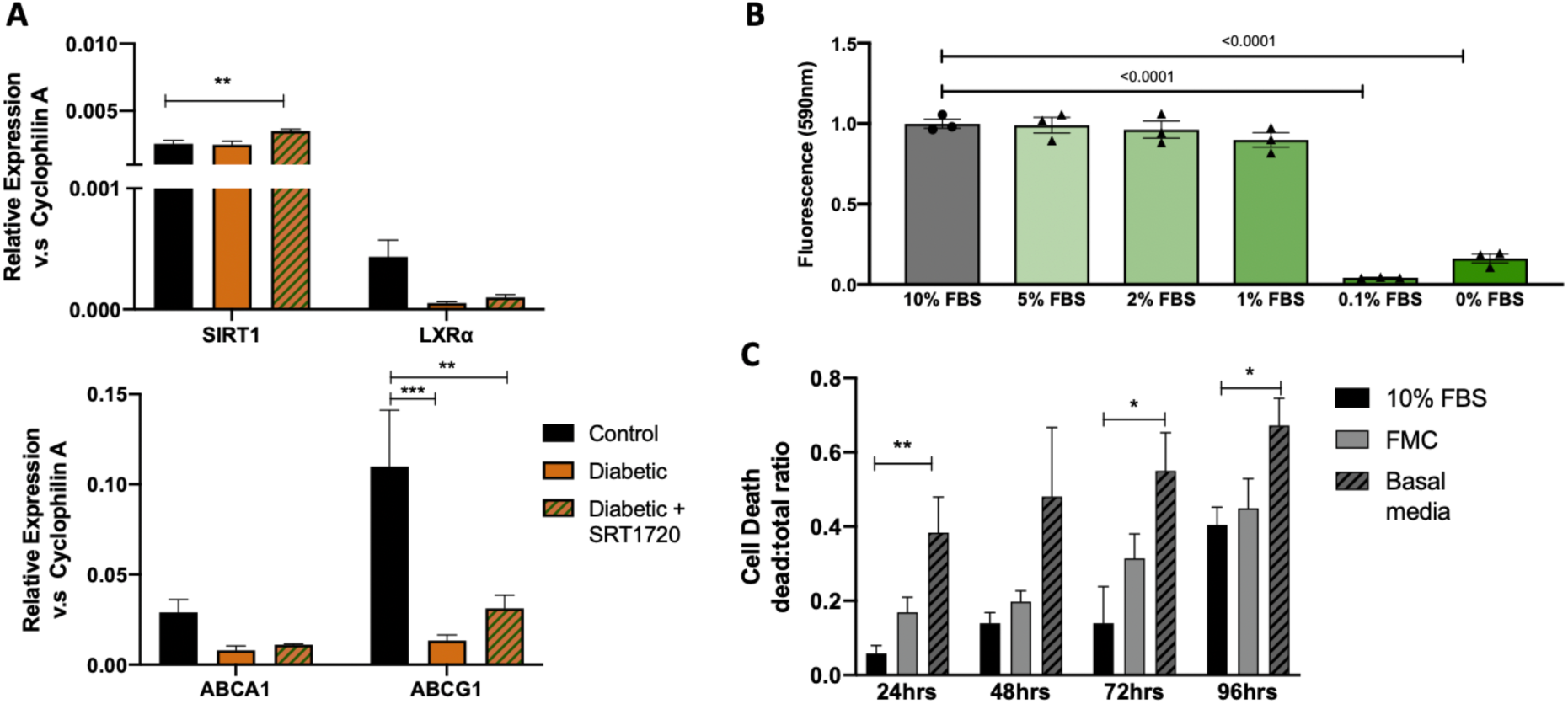
**A)** qRTPCR of SIRT1, LXRα, ABCA1 and ABCG1 in M2 macrophages from control, diabetic or diabetic + SRT1720 mice. n= 5 **B)** Cholesterol levels in bovine retinal endothelial cells (BREC) cultured in varying percentages of serum. n=3. **C)** Cell death (trypan blue exclusion assay). Basal media is BREC media without glucose, growth factors and other supplements found in the FMC culture media. n=3;*p<0.01, **p<0.001, ***p<0.0001. Data are represented as mean ± SEM.

**Supplemental Figure 2.**
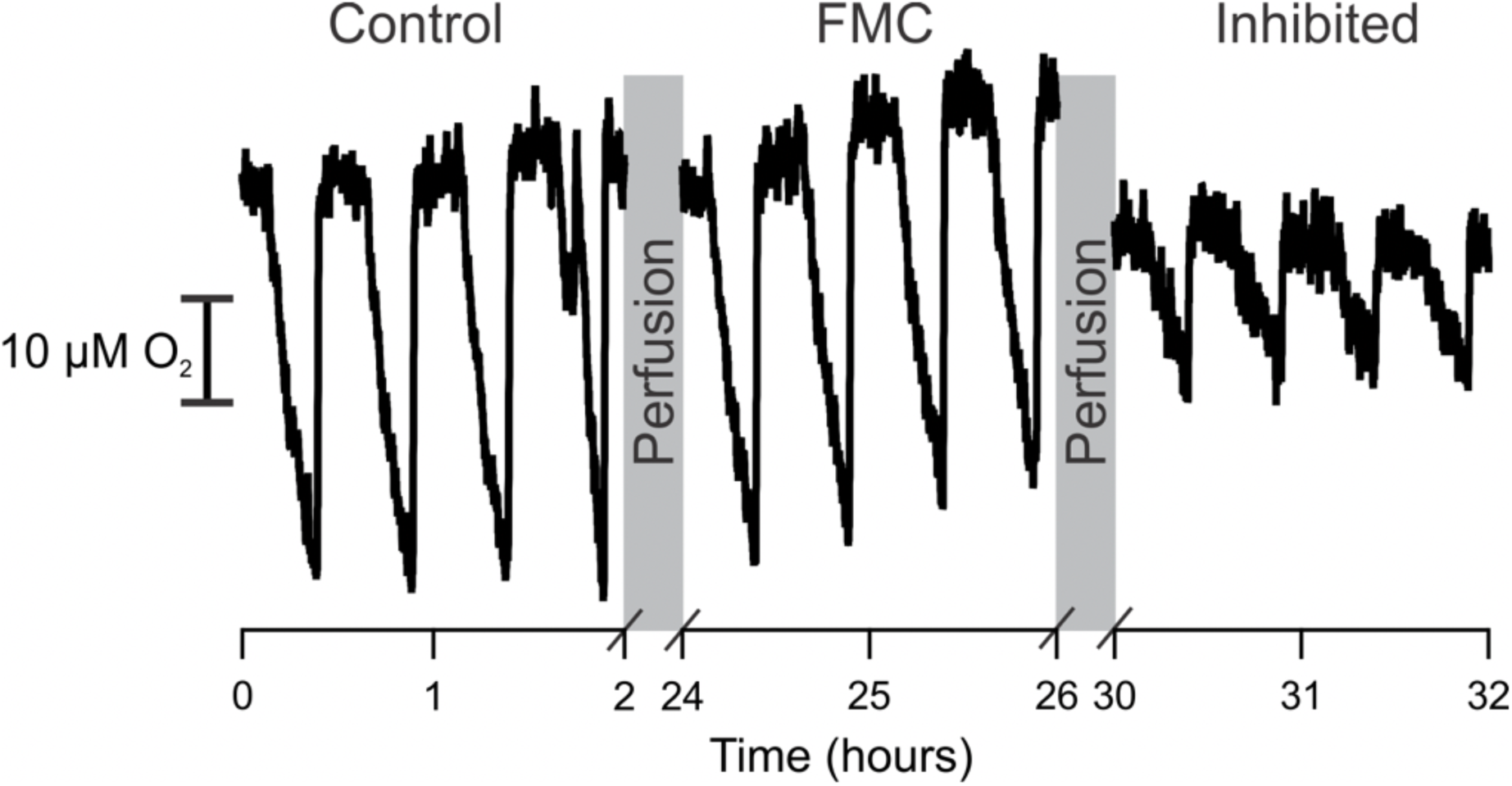
Endothelial cells were cultured on-chip and probed for substrate-supported respiration in the presence of serum (control) and 24 hours after removal of serum (FMC). Addition of potassium cyanide to the medium (inhibited) illustrates mitochondrial independent oxygen consumption. Bovine retinal endothelial cell respiratory activity does not change under FMC.

