## Supplemental Figure 1 for "Pharmacological and fasting-induced activation of SIRT1/LXRα signaling alleviates diabetes-induced retinopathy"

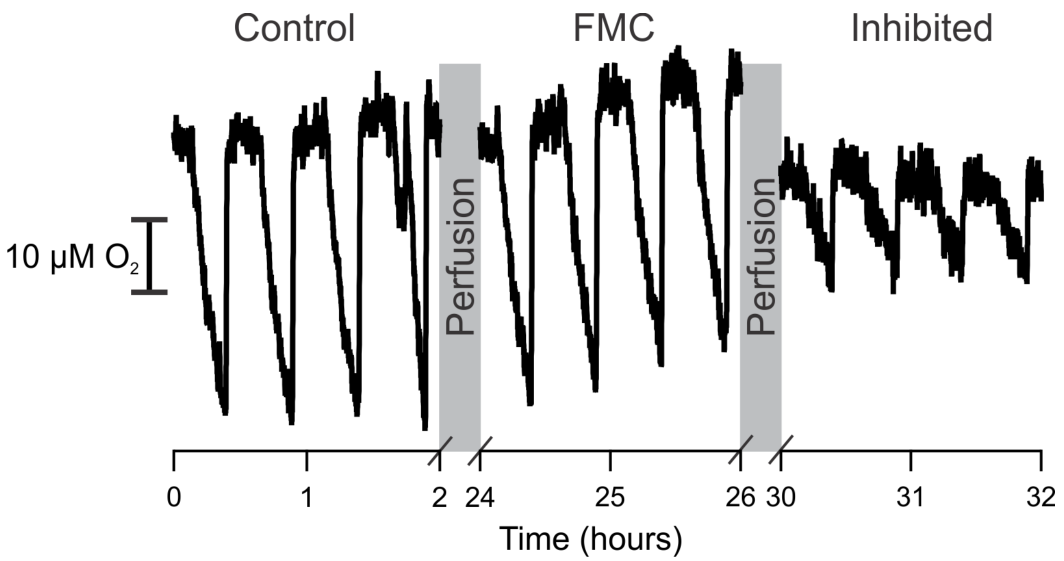


**Figure S2: Bovine retinal endothelial cell respiratory activity does not change under FMC.** Endothelial cells were cultured on-chip and probed for substrate-supported respiratory activity in the presence of serum (control) and 24 hours after removal of serum (FMC). Potassium cyanide (inhibited) illustrates mitochondrial independent oxygen consumption.
