## Supplemental Figure 2 for "Pharmacological and fasting-induced activation of SIRT1/LXRα signaling alleviates diabetes-induced retinopathy"

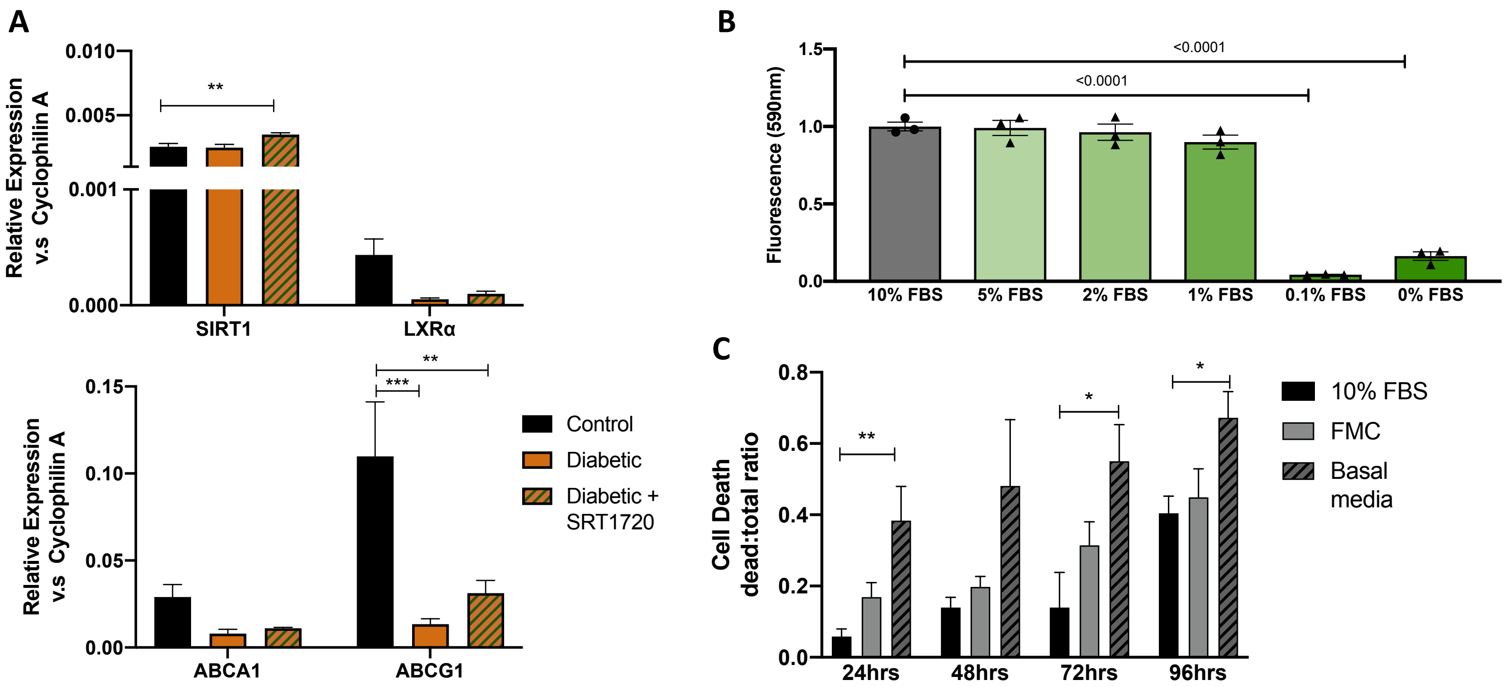


**Supplemental Figure 1. A)** qRTPCR of SIRT1, LXRα, ABCA1 and ABCG1 in M2 macrophages from control, diabetic or diabetic + SRT1720 mice. n= 5 **B)** Cholesterol levels in Bovine retinal endothelial cells (BREC) cultured in varying percentages of serum. n=3. **C)** Cell death (trypan blue exclusion assay). Basal media is BREC media without glucose, growth factors and other supplements found in the FMC culture media. n=3;*p<0.01, **p<0.001, ***p<0.0001.
